# Variation in root exudate composition influences soil microbiome membership and function

**DOI:** 10.1101/2021.11.04.467385

**Authors:** Valerie A Seitz, Bridget B McGivern, Mikayla A Borton, Jacqueline M Chaparro, Rebecca A Daly, Amy M Sheflin, Stephen Kresovich, Lindsay Shields, Meagan E Schipanski, Kelly C Wrighton, Jessica E Prenni

**Author notes:** Corresponding Author: J. Prenni, 1173 Campus Delivery, Colorado State University, Fort Collins, CO, 80523.

## Abstract

Root exudation is one of the primary processes that mediate interactions between plant roots, microorganisms, and the soil matrix. Previous research has shown that plant root exudate profiles vary between species and genotypes which can likely support different microbial associations. Here, utilizing distinct sorghum genotypes as a model system, we characterized the chemical heterogeneity between root exudates and the effects of that variability on soil microbial membership and metabolisms. Distinct exudate chemical profiles were quantified and used to formulate synthetic root exudate treatments, a High Organic acid Treatment (HOT) and a High Sugar Treatment (HST). Root exudate treatments were added to laboratory soil reactors and 16S rRNA gene profiling illustrated distinct microbial membership in response to HST or HOT amendments. Alpha and beta diversity metrics were significantly different between treatments, (Shannon’s, *p* < 0.0001, mrpp = 0.01, respectively). Exometabolite production was highest in the HST, with increased production of key organic acids, non-proteinogenic amino acids, and three plant growth-promoting phytohormones (benzoic acid, salicylic acid, indole-3-acetic acid), suggesting plant-derived sugars fuel microbial carbon metabolism and contribute to phytohormone production. Linking the metabolic capacity of metagenome-assembled genomes in the HST to the exometabolite patterns, we identified potential plant growth-promoting microorganisms that could produce these phytohormones. Our findings emphasize the tractability of high-resolution multi-omics tools to investigate soil microbiomes, opening the possibility of manipulating native microbial communities to improve specific soil microbial functions and enhance crop production.

**Importance:** Understanding interactions between plant root exudates and the soil microbiome provides an avenue for a more comprehensive appreciation for how plant roots modulate their microbial counterparts to promote an environment favorable to plant fitness. Although these dynamics are appreciated as indispensable, mechanisms controlling specific rhizobiome membership and complexity are not fully understood. In this study, we investigate how variability in root exudation, modeled after differences observed between distinct sorghum genotypes, contributes to altered microbial membership and metabolisms. The results demonstrate how microbial diversity is influenced by root exudates of differing chemical composition and how changes in microbial membership correspond to modifications in carbon utilization and enhance production of plant-relevant metabolites. Our findings suggest carbon substrate preferences among bacteria in semi-arid climate soils and mechanisms for root exudate utilization. These findings provide new information on plant-soil environments useful for the development of efficient and precise microbiota management strategies in agricultural systems.

## Introduction

The dynamic region of plant root-soil interfaces, known as the rhizobiome, is one of the most intricate ecosystems on earth, where complex interactions between plant roots, bacteria, fungi, and viruses occur (1, 2). The rhizobiome is under constant influence of varying biotic and abiotic factors including soil niches driven by soil type, land management, geographical location, soil pH, soil aggregate stability, and soil moisture content (3-6). The soil microbial communities may also be influenced by a range of biotic factors primarily derived from the presence of plant roots, and these changes can vary based on plant type, both at the species and genotype level (7-11).

While these abiotic and biotic factors all play a critical role in shaping a rhizobiome suitable to support healthy crops and plants, perhaps most influential is the chemical microenvironments produced by the plant through the release of photosynthetically-derived compounds into the soil, a process known as root exudation (12-14). Root exudate compounds can range from small (e.g. carbohydrates, amino acids, organic acids, phytohormones) to large (e.g. proteins, mucilage) molecules and root exudate profiles have chemical signatures unique to any given plant species and genotype (15, 16). Estimates of these photosynthetically-derived carbon molecules exuded from plant roots range from 3-40% depending on plant type and age, and these compounds influence a process known as the rhizosphere effect (9, 17). The rhizosphere effect results from the stimulation of microbial activity, leading to the development of a distinct rhizobiome.

Plant root exudates provide microbial substrates, as well as signaling molecules, that stimulate microbial activity and modify the local soil biogeochemistry. Under positive plant-rhizobiome interactions, plant growth-promoting rhizobacteria (PGPR) promote plant development by producing plant growth-promoting hormones like indole-3-actic acid (IAA), gibberellins, cytokinins, abscisic acid, and ethylene, suggested to modulate host plant physiological health (18, 19). Apart from producing plant growth-promoting compounds, PGPR can respond to root exudation signals to suppress pathogens, fix atmospheric nitrogen, and produce metal chelators that increase metallic soil micronutrient availability, among other positive outcomes (12, 20-22). Hence, root exudates enable plants to cultivate a favorable environment for microbial counterparts, which in return trigger the production and scavenging of nutrients, synthesize growth-promoting hormones, and facilitate survival against abiotic and biotic factors. Despite the recognized importance of plant-microbe mutualisms, an existing knowledge gap includes how genotypic differences in root exudation can shape soil microbiomes, as prior studies focused only on profiling a small subset of plant root exudates (23-26). Furthermore, investigations reporting more chemically-thorough root exudation characterization have relied on lower resolution microbial methodologies (e.g. phospholipid fatty acid analysis; PLFA) (27). Therefore, there is a current need for studies that employ high-resolution methods for both microbial and chemical characterizations to provide more mechanistic descriptions of exudate-microbe interactions.

Here, we used a multi-omics approach to characterize how root exudation can drive soil microbial community structure and function. Using three common Sorghum (*Sorghum bicolor* (L.) Moench) genotypes as a model, we determined the individual root exudate chemical profiles and used these to design relevant exudate amendments fed to agricultural soil microcosms in the absence of plant roots. We hypothesized that genotypic variation in root exudate chemical composition would enrich for distinct microbial populations. We tracked these microcosms over 20 days using exometabolomics, 16S rRNA gene profiling, and genome-resolved metagenomics. Integrating these data, we show that different exudate treatments led to distinct microbial communities and metabolisms, including production of plant beneficial metabolites. Harnessing this knowledge could support the growing need for sustainable agroecosystems by developing holistic agricultural management strategies that optimize the metabolic capabilities of the soil microbiome.

## Results & Discussion

### Sorghum genetics influence root exudation patterns

Sorghum is one of the most widely produced agricultural crops in the world, serving as a grain, forage, and cover crop for use as human food, livestock feed, and as a biofuel feedstock (28, 29). Due to its diverse agronomic uses, we leveraged three sorghum genotypes known to have distinct aboveground phenotypes that we hypothesized would contribute to distinct root exudate profiles. We selected 1) the grain sorghum, ‘BTx623’; 2) the sweet sorghum type, Leoti; and 3) the bioenergy sorghum PI 505735 (29-31). We grew sorghum genotypes *in vitro* (hydroponically) for seven days, and soluble exudates were collected in water and analyzed with non-targeted gas chromatography-mass spectrometry (GC-MS).

Two out of three sorghum genotypes produced metabolically distinct root exudate profiles. Metabolites from sorghum seedlings spanned known root exudate metabolite classes ranging from sugars, sugar alcohols, organic acids, and amino acids (**Fig. 1A, Fig. S1**). The most distinct chemical differences were observed between genotypes BTx623 and Leoti (**Fig. 1A**), while PI 505735 lacked a unique root exudate chemical profile and thus was not used in subsequent experiments. Among BTx623 and Leoti, genotype was a significant source of variation across organic acid and sugar root exudate composition (ANOVA, *p* < 0.001). Leoti was significantly enriched in organic and amino acids, while BTx623 was enriched in monosaccharides and disaccharides (**Fig. 1A, File S1**). Our hydroponically-derived metabolite findings suggest that sorghum genotypes release root exudates with distinct metabolic profiles and as such, we sought to see if these two exudate regimes would differentially enrich soil microbiome membership or metabolisms.

**Figure 1.**
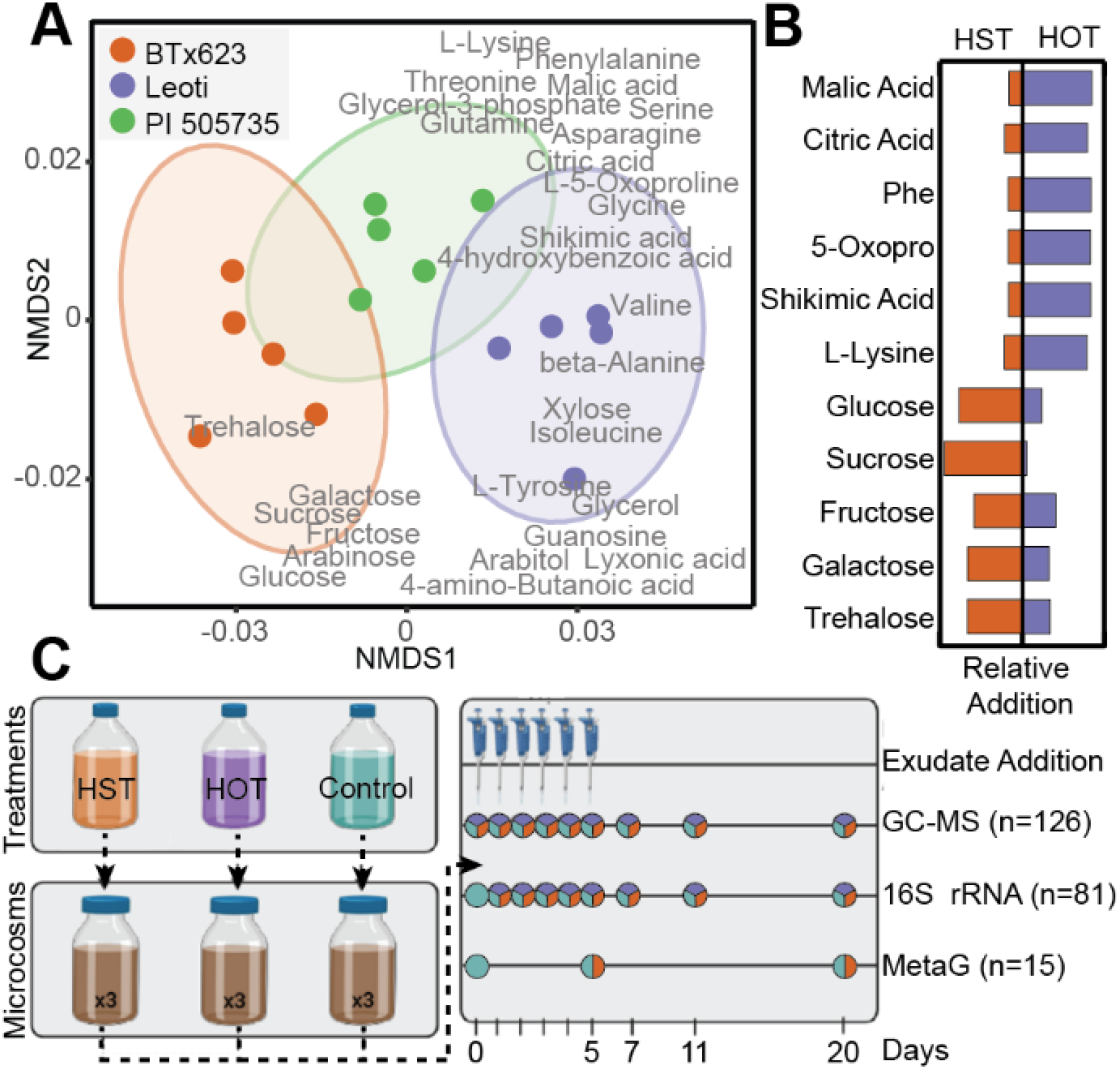
Root exudation metabolite profiles from three sorghum genotypes were used to design relevant amendments for soil microcosms. A) Non-metric multidimensional scaling (NMDS) of Bray-Curtis distances across metabolite abundance of Leoti (purple), BTx623 (orange), and PI 505735 (green) sorghum genotypes (stress=0.06). Ellipses denote the 95% confidence interval for each treatment. Significant metabolite loadings (p-value <0.05) are labelled in gray. B) Butterfly plots show the relative amount of each compound in the exudate additions for each treatment (Phe, Phenylalanine; 5-Oxopro, 5-Oxoproline); orange represents concentrations for High-Sugar Treatment (HST) and purple represents concentrations for the High Organic acid Treatment (HOT). C) Microcosm schematic depicts treatment formulation from where HOT represents Leoti, HST represents BTx623, and Control was buffered media lacking an exudate treatment. Triplicate microcosms with agricultural soil were maintained for a 20-day experiment with the sampling schematic denoting when samples were obtained. Time points with circles represent the samples taken each day, with the total number (n) of samples for each analysis listed. Colors within circles represent the type of sample (HST = orange, HOT = purple, control = teal) taken at that time point.

### Controlled laboratory microcosms for tracking root exudate influence on soil microbial communities

The two sorghum genotypes were used to formulate synthetic root exudate treatments for evaluating the impact of root exudate composition in the absence of roots on soil microbial community structure and function. Specifically, we formulated root exudate solutions for laboratory-scale soil reactors (microcosms) amended with either a high sugar treatment (HST) representing BTx623, a high organic acid treatment (HOT) representing Leoti, or exudate-lacking media control. Both HST- and HOT-amended microcosms received the same eleven root exudate compounds (glucose, galactose, fructose, sucrose, trehalose, malic acid, lysine, phenylalanine, 5-oxoproline, citric acid, and shikimic acid), but in varying concentrations to model the concentrations observed in BTx623 and Leoti exudates (**Fig. 1B**). The soil microcosms were constructed using soil from semi-arid agricultural plots and were amended with exudate treatments daily for 5 days. We hypothesized these exudate-informed amendments would enrich for distinct microbial communities at both a community and functional potential level. To test this, microcosms were tracked for chemical and microbiological analyses during and after the period of exudate addition (**Fig. 1C**), offering a new opportunity to uncover the impact of root exudates on soil microbial communities.

### High sugar and high organic acid exudate treatments structure agricultural soil microbial communities

We first used 16S rRNA gene amplicon sequencing to temporally profile the microbial diversity and membership across our three treatments (HOT, HST, exudate control, **Fig. 1C**). In total, we sequenced 81 samples, generating 2,148,831 high-quality reads with an average of more than 30,000 reads per sample (**File S2**). After denoising, a total of 9,818 amplicon sequencing variants (ASVs) were detected, representing 43 phyla (**File S2**). We first assessed alpha and beta diversity metrics to understand soil microbial community changes in response to exudates. Over time, we observed distinct changes in response to amendments with either the HST or HOT (**Fig. 2A**). For instance, after one day of exudate amendment, HST-amended microcosms saw significant decreases in ASV species richness (*p* = 0.007), Shannon’s Diversity Index (*p* = 0.019), and Pielou’s evenness (*p* = 0.017). In contrast, we did not detect significant changes in these metrics for HOT or control microcosms in this time period. This suggests HST-amendment strongly enriches for select members of the microbial community. In support of this, we observed an ASV belonging to the genus *Psuedomonas* (**File S2**) accounting for 59.4% of the day 1 community relative abundance in HST microcosms. Following 5 days of exudate addition, the HST-amended microcosms retained low Shannon’s (*p* = 0.038), low ASV richness (*p* = 0.042), and evenness (*p* = 0.017) compared to the HOT and control microcosms (**Fig. 2A)**. Interestingly, after amendments were stopped, these diversity metrics were no longer significantly different between treatments, highlighting the importance of these amendments in structuring the microbial communities.

**Figure 2.**
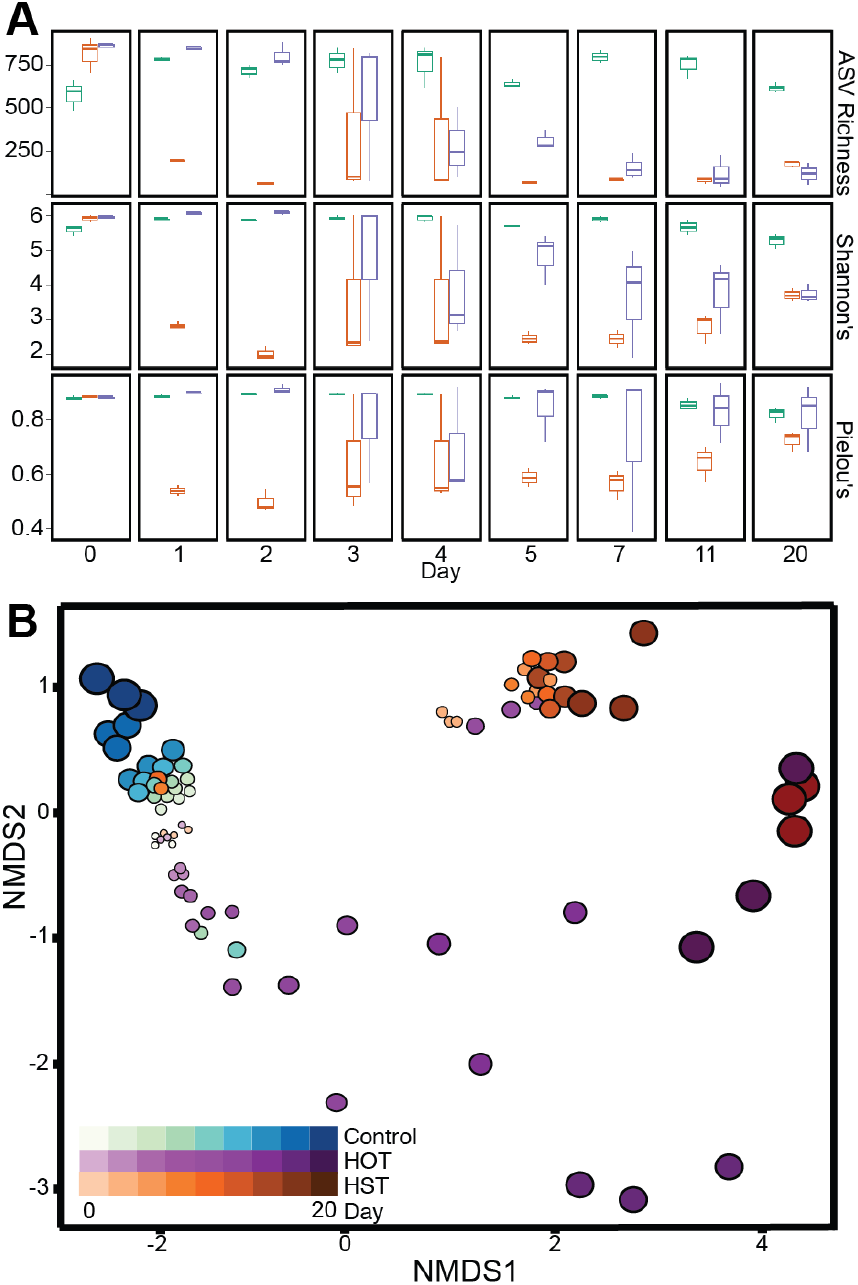
16S rRNA gene diversity metrics and membership changes with exudate amendment. A) ASV richness (top), Shannon’s Diversity Index (middle), and Pielou’s Evenness for each day highlighting differences in treatment richness, alpha diversity, and community evenness. Coloring corresponds to treatment: control (teal), HST (orange), and HOT (purple). B) Non-metric multidimensional scaling (NMDS) of Bray-Curtis distances of 16S rRNA amplicon communities showing changes in microbial community structure and membership over time with colors representing treatment and the size and darkness of circles representing time.

Next, the microbial community similarity across treatment was assessed using Bray Curtis dissimilarity matrix and visualized using non-metric multidimensional scaling (NMDS) (**Fig. 2B**). We found that HST and HOT amendments significantly shifted microbial community composition and structure during and after periods of synthetic root exudate addition (multi-response permutation procedure, mrpp = 0.01, p-value < 0.05). While the control exudate communities remained stable over time, we observed distinct temporal shifts in both HOT and HST microbial communities by day 20 (**Fig. 2B**). To quantify differences across treatments, we used beta dispersion analysis with ANOVA to visualize group dispersions and test significance of differential compositions due to exudate amendments. This analysis revealed that microbial communities were significantly altered across treatments (*p* < 0.0001) and time (*p* < 0.0001). These microbial community alpha and beta diversity metrics highlight that both treatment and time were factors shaping the soil microbiome in both the HST- and HOT-amended microcosms. Scaling beyond these microcosms, this highlights the need for understanding the composition and temporal dynamics of natural root exudates to develop exudation-focused crop management strategies.

### Exometabolites hint at potential microbial metabolisms stimulated by exudates

In these soil reactors, we tracked the temporal dynamics of exometabolites (**Fig. 1C**), which are the extracellular fraction of molecules that are inferred to be produced and or utilized by soil microorganisms. We classified and assigned the detected exometabolites to three chemical classes: (i) central carbon metabolism, (ii) amino acids and derived compounds, and (iii) phytohormones (**File S1)**. The exometabolites were coordinated to the microbial communities identified by 16S rRNA gene sequencing (**Fig. S2**) and exhibited differences by treatment (**Fig. 3**). Relative to the HOT and controls, the HST was enriched in most of the detected exometabolites, likely reflecting a greater metabolic stimulation from the sugar rich treatment (**Fig. 3**). This finding is not completely unexpected, as prior studies have noted sugar metabolism is more efficient than amino acid metabolism in soil microorganisms (32). Metabolites relevant to microbial central carbon metabolism were some of the most enriched exometabolites in HST microcosms (**Fig. 3A**). We detected six organic acids that changed significantly over time and were most dynamic in HST. For example, between days 2 and 3, succinic acid and fumaric acid increased 1.5 and 0.9-fold (log2), respectively, in the HST-amended microcosms relative to HOT and the control treatments (**Fig. 3A)**. We also detected increases in itaconic acid (0.7-fold) and oxalic acid (1.6-fold) over time, both of which can be derived from tricarboxylic acid (TCA) cycle intermediates cis-aconitate and oxaloacetate, respectively (33). Furthermore, consumption of malate and citrate was inferred in these microcosms at the same time points due to their loss over time (**Fig. S3**). Perhaps indicating metabolite cross feeding across the community, we also observed a spike in pyruvate at days 2-3 in the HST, followed by consumption concomitant with the production of lactic acid across the experiment (**Fig. 3A)**. Collectively, this exometabolite data pointed to HST exudate treatments differentially altering soil microbial central carbon metabolism, illustrating important impacts for microbial cross feeding and carbon metabolism in the soil.

**Figure 3.**
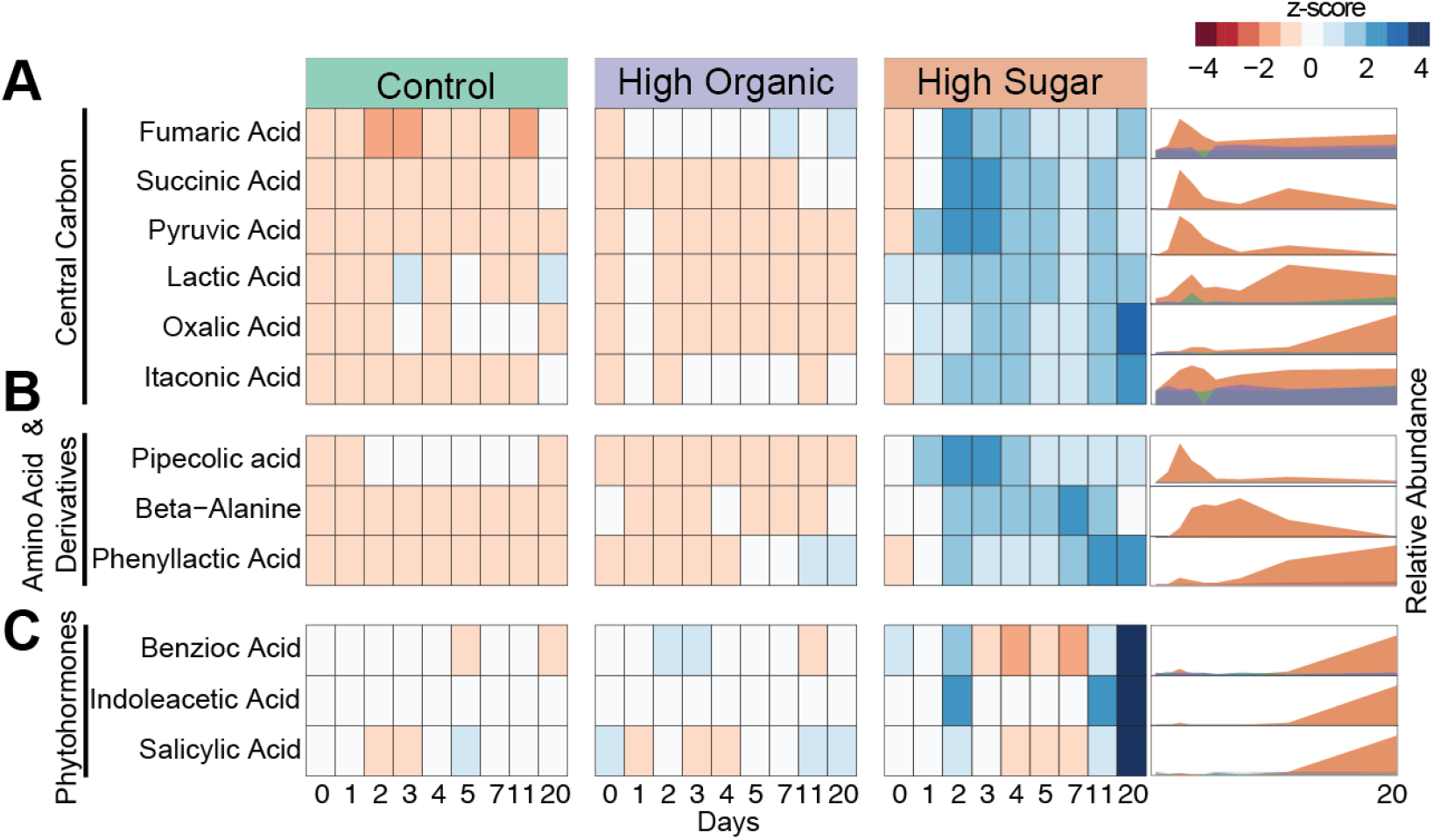
Exometabolite abundances across treatments. Heatmaps (left) showing the relative abundances (represented as a z-score across samples) of each exometabolite detected. Time increases from left (day 0) to right (day 20) within each treatment. Ridgeline plots (right) show the relative abundance of these exometabolites over time. Coloring corresponds to treatment: control (teal), HST (orange), and HOT (purple).

We also detected chemical evidence for production and consumption of amino acid and amino acid derived compounds by these soil microbial communities. Similar to central carbon metabolites, these organic nitrogen exometabolites were most enriched in HST relative to the other treatments, a somewhat unexpected response given the HOT amendments were initially dosed with these types of compounds (**Fig. 1B**). We observed the production of two non-proteinogenic amino acids (β-alanine and pipecolic acid) and one aromatic compound (phenyllactic acid) (**Fig. 3B)**. β-alanine, which was produced over the first 7 days but degraded by day 20, is an important amino acid precursor synthesized from L-aspartate and is necessary for biosynthesis of coenzyme A from the vitamin pantothenate (34). Next, we observed the amino acid, pipecolic acid (PA), whose production peaks between days 2-4 in HST-amended microcosms and is inferred to be microbially synthesized from lysine (35). In support of this, lysine, part of the exudate treatments, was consumed during this period (**Fig. S3)**. PA synthesis has broader ramifications for the entire microbiome, as it is a required intermediate in microbial secondary metabolite production of antibiotics and anthelmintics (35). PA dynamics, which indicate production and subsequent consumption by day 20, suggest its use as a public good (36). Finally, phenyllactic acid (PLA), a phenylalanine derivative, increased 2.6-fold in the HST microcosms over time, peaking at day 11 and 20 (**Fig. 3B**). Microbes that release amino acid and amino acid derived compounds could be competitive root colonizers or act as antagonistic agents against target pathogens (37).

Most notably, we observed a significant increase in the abundance of three phytohormones between days 0 and 20 (**Fig. 3C)**. Salicylic acid (SA), benzoic acid (BA), and indole-3-acetic acid (IAA) significantly increased (*p* < 0.01) from days 0 to 20 in the HST relative to microcosms treated with HOT or exudate-lacking control. In HST microcosms, BA and SA increased 1-fold (log2) while IAA increased the most with a 3-fold (log2) increase from day 0 to day 20. In plants, SA, BA, and IAA are vital phytohormones integral to physiological processes like plant defense and development (38), and these compounds have been suggested to mediate symbiotic plant-microbe relationships in crop plants such as oat and corn (13, 39). Our detection of these metabolites over time highlights that certain root exudation chemical profiles can stimulate soil microbes to produce phytohormones critical for plant growth and defense.

### Curation of a semi-arid climate agricultural soil MAG database

In light of the phytohormone production observed in our HST microcosms, we sought to identify microbial genomes capable of producing these compounds. Towards this goal, we constructed a database of semi-arid, agricultural soil metagenome-assembled genomes (MAGs) representing the suite of soil microbes in the HST and control microcosms. To maximize MAG recovery, we obtained more than 365 Gigabasepairs total of sequencing from control and HST microcosms at three different metabolically-relevant timepoints (control: day 0, 5, 20; HST: day 5, 20, each in triplicate; n= 15 metagenomes, **Fig. 1C**). With this data, we reconstructed 371 MAGs that were dereplicated at 99% identity into 243 MAGs, of which 28% were high-quality (40) (**Fig. 4A, File S2**).

**Figure 4.**
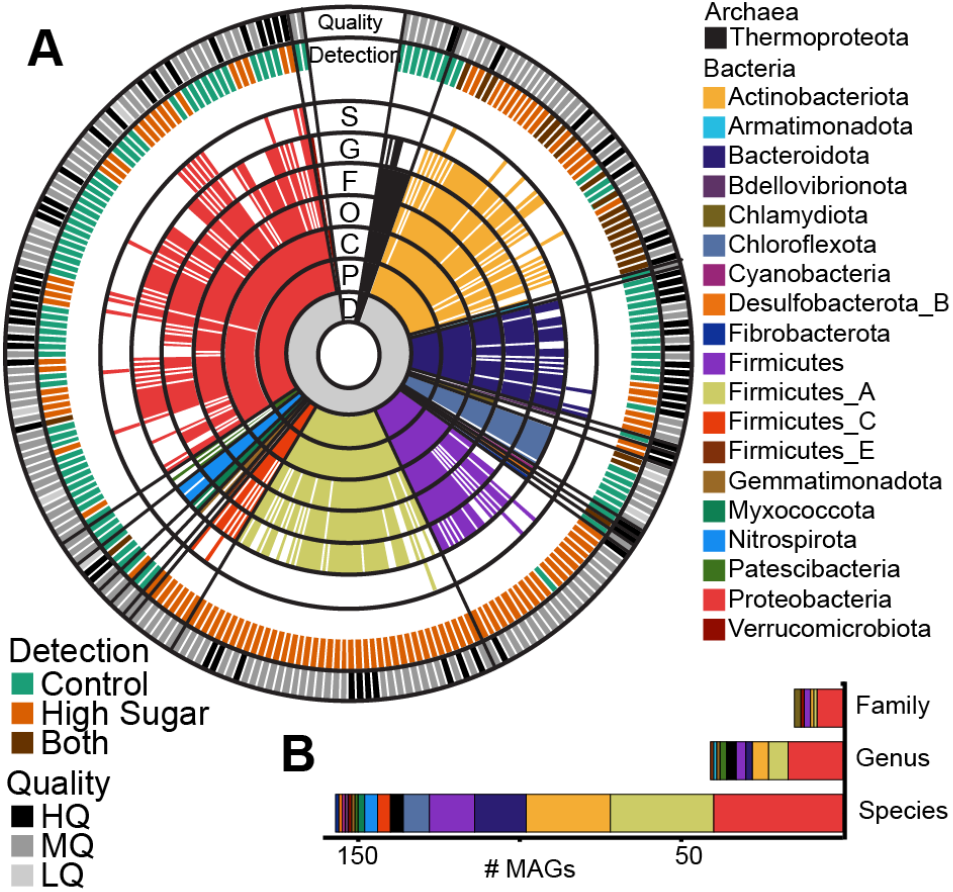
Taxonomy of the 243 dereplicated metagenome assembled genomes in the genome database. A) Sequential colored rings indicate the most resolved taxonomic level that could be assigned by GTDB-tk (70). Taxonomic level (D=Domain, P=Phylum, C=Class, O=Order, F=Family, G=Genus, S=Species) is denoted in black with a single letter abbreviation. Ring color corresponds to phylum assignment, with the color legend at right. The treatment condition corresponding to MAG detection is illustrated in the outer ring labeled “Detection”; MAGs detected only in Control metagenomes are indicated by teal, MAGs detected only in HST metagenomes are indicated by orange, and MAGs that were detected in both indicated with brown (see Experimental Procedures for detection thresholds). The MAG quality is shown in the outermost ring following the MIMAG standards (40): high quality (HQ, >90% complete, <5% contamination), medium quality (MQ, >50% complete, <10% contamination), and low quality (LQ, here defined as >48% complete, <10% contamination). B) Stacked bar graph shows the number of dereplicated MAGs recovered that represent novel families, genera, or species according to taxonomy assignments from GTDB-tk. Coloring corresponds to MAG phylum.

Of these MAGs, 49% (n=119) were detected in HST-amended microcosms while 41% (n=100) were detected in the control microcosms at any time point (**Fig. 4A, File S2**). Notably, only 10% (n=24) of these MAGs were shared across the two treatments at any one timepoint, further supporting our 16S rRNA and metabolite data showing HST amendments significantly altered the soil microbial community. Our MAG database contained representative soil microbes across 20 phyla, including MAGs belonging to 15 previously unidentified families (6%), 41 previously unidentified genera (17%), and 157 previously unidentified species (65%) (**Fig. 4B**). We recovered 26 MAGs with partial or full 16S rRNA genes and could directly link 19 of these MAGs to 16S rRNA amplicon sequencing identified ASVs, providing metabolic blueprints for these taxa. This highlights the advantage of coupling community profiling (16S rRNA gene) with high resolution metagenomics to capture the functional potential encoded within the soil microbiome.

### Biosynthesis of three phytohormones represent potential plant growth promoting rhizobacteria (PGPR) within MAG database

#### Bacterial salicylic acid production is assigned to a new species of *Pseudomonas*

Our multi-method approach suggested linkages between root exudate-induced microbiome community shifts and specific metabolic functions, with implications for phytohormone producing bacteria. To assign metabolic roles to specific organisms for phytohormone production, we examined MAG metabolic potential for salicylic acid, benzoic acid, and indoleacetic acid.

Salicylic acid is synthesized by plants in response to abiotic stressors (e.g. salinity, cold, drought stress) or for developmental signaling cascades for flowering and senescence (41). It is known that PGPR can also produce SA, representing a beneficial service to the plant host (39, 42, 43). In our MAGs, we tracked SA biosynthesis from the conversion of chorismate to isochorismate via isochorismate synthase (*pchA*), and subsequent conversion to salicylate by isochorismate pyruvate lyase (*pchB*) (**Fig. 5A**) (44). From our MAG database, two HST-detected MAGs encoded a *pchA* and one MAG encoded a homolog for *pchB*. Notably, a single MAG representing a novel species within the genus *Pseudomonas_E* (L_E1_T20_B_bin.65) encoded two copies of *pchAB* for SA biosynthesis via the chorismate pathway (**Fig. 5B**). Linking the observed increase in SA in our exometabolite data with the potential organism capable of completing this pathway, this *Pseudomonas_E* MAG was highly abundant in HST metagenomes at day 20 (**Fig. 5E**), mirroring when SA was produced. Taken together, these results present *Pseudomonas_E* as the most likely candidate for SA production in high sugar treated microcosms (**Fig. 5C**).

**Figure 5.**
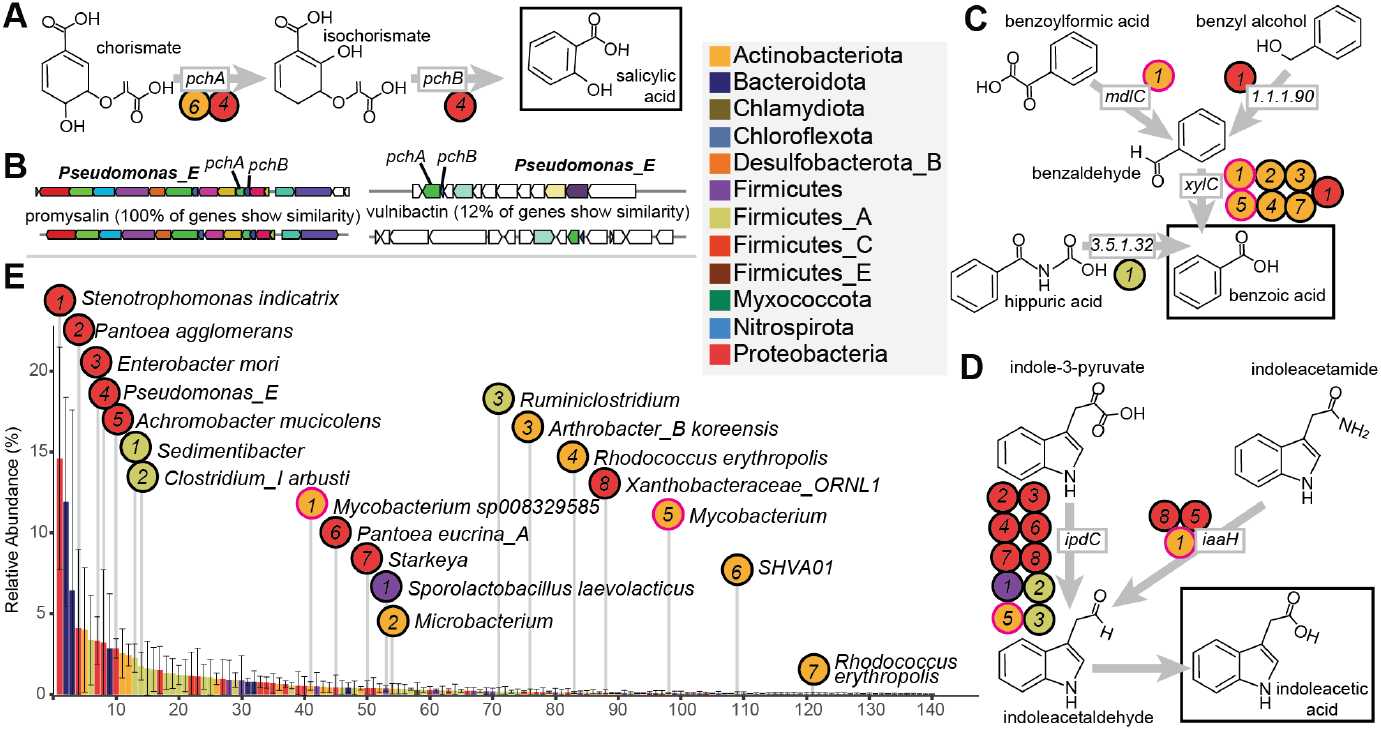
Diverse MAGs encode biosynthetic potential for salicylic acid, benzoic acid, and indoleacetic acid. A) Salicylic acid production from chorismate by *pchAB*. B) A *Pseudomonas* MAG encoded two biosynthetic gene clusters containing *pchAB*, including one for the antimicrobial Promysalin (left) and another for a predicted siderophore (right). The clusters on the bottom are the reference clusters from antiSMASH. C) Benzoic acid production from multiple pathways. D) Indoleacetic acid production pathways. E) Rank abundance curve of MAGs detected in HST metagenomes at day 20. Bars represent the average relative abundance (n=3), and error bars represent one standard deviation. Bars are colored by MAG phylum, and key phytohormone producing MAGs are indicated. In *A, C*, and *D*, circles correspond to MAGs encoding each gene in E. The circles for the two *Mycobacterium* MAGs with BA and IAA potential are outlined in pink.

Previous research investigating the context for bacterial salicylate biosynthesis is limited, however, some research has suggested this compound is used as a siderophore (42) or can support antibiotic production (45). Excitingly, analysis of the *Pseudomonas_*E MAG revealed one set of *pchAB* which occurred in a predicted siderophore biosynthetic gene cluster (**Fig. 5B**). This could support that SA production is an iron acquisition strategy in the absence of plant roots, as microbes rely on plant metal chelators in iron-limited soils, which is a common condition of rhizospheric soils (46). Furthermore, the second *pchAB* cluster occurred in a 16kb gene cluster that showed 100% gene similarity to the promysalin biosynthetic gene cluster from *Pseudomonas putida* (**Fig. 5B**). Promysalin is an antibiotic that contains SA and was shown to be selectively antagonistic to other closely related pseudomonads and suggested to be involved in rhizosphere colonization (45). Together, our integrated MAG and exometabolite data suggest HST created conditions that stimulated the *Pseudomans_E* MAG to produce SA, which could ultimately aid in its competition for limited resources in the complex soil microbiome.

#### Diverse benzoic acid production pathways encoded by multiple MAGs

Benzoic acid (BA) is known to improve stress tolerance and contribute to growth regulators in plants (47). BA can be microbially-produced through the degradation of several aromatic compounds (**Fig. 5C**). The terminal step of most of these pathways is the oxidation of benzaldehyde to BA by benzaldehyde dehydrogenase (*xylC*). Seven MAGs encoded *xylC*, including *Stenotrophomonas indicatrix, Arthrobacter koreensis*, two MAGs of *Rhodococcus erythropolis*, and two species of *Mycobacterium* (**Fig. 5C**). These MAGs were only detected in the HST treatment at day 20 (**Fig. 5E**), matching when BA was observed in exometabolomic data (**Fig. 3C**).

MAG analysis revealed the *S. indicatrix* MAG encoded an aryl-alcohol dehydrogenase (EC: 1.1.1.90) adjacent to *xylC*, which would enable reduction of benzyl alcohol to benzaldehyde (**Fig. 5C**). This MAG is also the most abundant MAG in HST metagenomes at day 20 (**Fig. 5E**), indicating it may be important to BA production in the microcosms. Beyond *S. indicatrix*, a MAG representing *Mycobacterium* sp008329585 encoded a benzoylformate decarboxylase (*mdlC*) which decarboxylates benzoylformic acid to benzaldehyde (**Fig. 5C**). Finally, an abundant MAG representing a novel species of *Sedimentibacter* encoded a gene for hippurate hydrolase (**Fig. 5E**). This enzyme cleaves glycine from hippuric acid, producing BA (EC: 3.5.1.32, **Fig. 5C)**. In all, our MAG-resolved analysis revealed BA may be produced by several MAGs through a variety of pathways, illustrating the metabolic versatility harbored in the soil microbiome.

#### Bacterial indoleacetic acid production encoded by diverse MAGs

Indoleacetic acid is a phytohormone in the auxin family and is important for plant growth regulation including enhanced growth, increased root biomass, and proper development (48, 49). From a microbial perspective, IAA production offers a competitive colonization strategy over non-IAA producing strains, and also stimulates more exudate production, providing a carbon rich unique niche to support IAA synthesizers (50). Tryptophan (Trp) is the primary precursor for IAA biosynthesis in microorganisms (51), with five known biosynthetic pathways characterized (52). We surveyed our MAG database for these genes and found 14 HST-detected MAGs capable of IAA production using two different pathways.

Nine MAGs encoded indole-pyruvate decarboxylase (*ipdC*), the key gene for converting Trp to IAA via indole-3-pyruvate (**Fig. 5D**). These MAGs spanned four phyla, with six belonging to the Proteobacteria, covering several of the most abundant MAGs at day 20 in the HST metagenomes (**Fig. 5E**). Beyond this pathway, three MAGs encoded *iaaH*, the key enzyme for IAA production via indoleacetamide (**Fig. 5D**). One of these MAGs, a novel species of *Xanthobacteraceae*, encoded both *iaaH* and *ipdC*. This fits with previous observations of single organisms encoding redundant IAA biosynthesis pathways (53). Of note, two Mycobacterial MAGs with IAA-production potential also had BA-production potential. One novel *Mycobacterium* species encoded *ipdC* and *xylC*, while *Mycobacterium* sp008329585 encoded *iaaH* in addition to *mdlC* and *xylC*, indicating some soil microbes can produce multiple phytohormones.

Collectively, our MAG-resolved survey of phytohormone production highlights several HST-enriched MAGs encode the capacity for SA, BA, and IAA production. Phytohormone production potential is encoded across both dominant and less abundant MAGs (**Fig. 5E**), highlighting the redundancy of this metabolism in the soil microbiome. It is important to note that the metagenomic results presented here only represent the genomic capabilities of a hypothesized genome, thus, additional experimental evaluation of gene expression would be required to definitively confirm specific bacterial synthesis of SA, BA, and IAA. Yet, we find these MAG results, coupled with exometabolite evidence for these compounds through time, to be an exciting platform for targeted studies aimed at harnessing the power of microbial metabolisms for improving agroecosystems.

## Conclusion

The majority of previous research investigating soil microbial responses to root exudates have largely only considered 16S rRNA gene profiling or PLFA to survey microbial behavior. This study gains a more comprehensive appreciation for the influence of plant root exudation on the soil microbiome by utilizing a suite of high-resolution multi-omics approaches to highlight microbial metabolisms that may influence rhizobiome function and overall plant health. We first showed that naturally derived root exudate amendments differentially impacted overall microbial community diversity. Second, shifts in microbial community structure mirrored differences in exometabolites and highlighted production of interesting plant-relevant metabolites. Finally, we linked exometabolite data to bacterial genomes by highlighting MAGs capable of phytohormone biosynthesis. Our findings resolve mechanisms by which root exudate composition influenced the functionality of the rhizobiome, uncovering possible important ramifications on the microbiome (e.g. metal acquisition, antibiotic production) and host plant physiology. Importantly, our MAG database can be leveraged as a genomic resource for agricultural soil microbiomes for other researchers working in semi-arid systems. Our findings emphasize the dynamic nature of plant-soil environments that could be leveraged to develop microbiota management strategies for improved soil microbial community function and enhanced crop production.

## Materials & Methods

### Sorghum Root Exudate Collections

Mature sorghum seeds for three sorghum genotypes (Leoti, BTx623, and PI 505735) were first sterilized by placing seeds in sterile 50 mL conical tubes with 45 mL 95% ethanol. Tubes were vortexed for 2 minutes and ethanol was removed. 45 mL of Captan fungicide solution (0.2g Captan fungicide in 45 mL sterile water) was added to remove any fungal components on the seeds and vortexed for 3 hours. After removal of the Captan fungicide, 45 mL of 100% Clorox bleach (8.25% sodium hypochlorite active ingredient) + 3 droplets of dish soap (Ajax Triple Action Orange) was added for the final sterilization step and seeds were shaken for 20 mins. In a sterile tissue culture hood, the bleach was removed using sterile techniques and the seeds were rinsed five times with sterile water. Seeds were blotted dry and placed on germination paper. The germination paper with the seeds was placed in a clean 600 mL beaker with a solution of 1 mM CaCl_2_. After 7 days of growth, the sorghum seedlings were removed from the paper and 30 seedlings were pooled into a 250 mL glass bottle filled with 80 mL of ultrapure water. Root exudates were collected for each genotype in separate bottles. The bottles were covered with aluminum foil to protect roots from the light and were placed on a rotary shaker. After 2 hours, the roots were removed, blotted dry, and weighed. The root exudate suspensions containing the root exudates were filtered through a 0.2 μm filter membrane to remove root detritus and microbial cells. Samples were frozen at -80°C before lyophilizing to sublimate any water. We recognize that the use of hydroponic growth chambers does not exactly replicate soil growth conditions of sorghum (54); however, this system provided a highly controllable, tractable, and sterile environment that eliminated confounding microbial or soil influences for more accurate downstream analytical detection.

Samples were lyophilized completely before resuspension in 15 mL sterile HPLC-grade water. Resuspended exudates were vortexed thoroughly to remove all residue from the bottle, transferred to a clean 50 mL falcon tube and dried completely under nitrogen gas. 1mL LCMS-grade 80% acetonitrile was added to the falcon tube with dried exudates and vortexed until thoroughly resuspended. 0.5 mL was transferred to a clean 2 mL glass vial, and 5 μg of glucose-^13^C_6_, D-arabinose-^13^C_5_, sucrose-^13^C_12_, galactose-^13^C_6,_and fructose-^13^C_6_ were added to each vial and dried completely under nitrogen prior to derivatization.

### Sample Derivatization for Sorghum Root Exudates

Dried samples were resuspended in 75 μL of 0.2M of methoxyamine hydrochloride in pyridine (Sigma), incubated at 60°C for 45 min, vortexed, sonicated for 10 min, incubated a second time at 60 °C for 45 min and allowed to cool to ambient temperature. 30 μL of each sample were then transferred to a separate 2mL glass vial for MSTFA derivatization and the other 45 μL for Acetic Anhydride (AA) derivatization. 90 μL of 100% AA was added to AA vials, vortexed for 30 s, incubated at 60°C for 60 min, and cooled for 10 min at ambient temperature. 30 μL of N-methyl-N-trimethylsilyltrifluoroacetamide plus 1% trimethylchlorosilane (MSTFA + 1% TMCS, Thermo Scientific) was added to the MSTFA vials, vortexed for 30 s, incubated at 60°C for 35 min and cooled to ambient temperature before transferring to vial inserts. AA samples were dried completely under nitrogen and resuspended in 40 μL of 100% ethyl acetate and then loaded into glass inserts for analysis.

### Non-Targeted and Targeted GC-MS Analysis for Sorghum Root Exudates

Metabolites were analyzed and detected using a Trace 1310 GC (Thermo) coupled to an ISQ mass spectrometer (Thermo). Samples (1 μL) were injected into an injection port at 285°C and 1:10 split ratio. Separation was accomplished with a 30 m TG-5MS column (Thermo Scientific, 0.25 mm i.d., 0.25 μm film thickness) and a helium gas at 1.2 mL/min flow rate. The oven temperature program started at 80°C for 30 sec, ramped to 330°C at 15°C/min, and then held at the final temperature for 8 min. The transfer line and ion source were maintained at 300°C and 260°C, respectively. Masses between 50-650 m/z were scanned at 5 scans/sec after electron impact ionization. A pooled QC sample (made by combining an equal volume of all samples) was injected after every 6 samples to monitor instrument stability throughout the analysis.

### Synthetic Exudate Preparation

Exudate treatments were formulated based on the results of the exudate profiles for the two most divergent sorghum genotypes (Leoti and BTx623). High Sugar Treatment (HST) was formulated to mimic BTx623 with higher sugar and lower organic acid composition and the High Organic acid Treatment (HOT) was formulated to mimic the Leoti exudate profiles with higher organic acid, and lower sugar composition (**Table S3**). The media control treatment (Control) was a 10mM phosphate buffer, consisting of ammonium chloride, disodium phosphate, and sodium dihydrogen at a pH of 6.5 (**Table S1**). Exudates found in each genotype were weighed out to their respective masses (**Table S3**) to supply equal exudates for 6 days of exudate addition to the microcosms, homogenized, suspended in 10mL 10mM phosphate buffer, vortexed, aliquoted into 6 tubes for each day of exudate addition, and frozen at -80°C until use. Aliquots were removed 30 min before use, incubated at 24°C until thawed, and added to microcosms at the respective time point.

### Soil Samples

The soil (microbial inoculum) was collected from agricultural fields at the Colorado State University Agricultural Research and Education Center (CSU-ARDEC) near Fort Collins, CO on October 4^th^, 2019. The climate at the site is semi-arid, with 408 mm mean annual precipitation and a mean annual temperature of 10.2°C (1981-2010 average, https://usclimatedata.com/). The soil is classified as an Aridic Haplustalf. Three 2-cm diameter soil core samples to approximately 15cm depth were collected from each of seven different plots. The soil was stored at -20°C until microcosm construction. 20g of soil from each replicate/plot was pooled and homogenized to create a representative bulk soil repository used in the following microcosms experiment.

### Microcosm Experimental Set-Up

Microcosms were established and sampled as previously described (55, 56). Briefly, 5 g of homogenized soil and 35mL of phosphate buffered (pH 6.5) minimal medium (**Table S1**) was added to sterile 50 mL serum vials to construct each microcosm. Microcosms were vortexed and allowed to settle for 5 min. Then day 0 samples were taken by removing 1 mL of soil slurry for exometabolomics analysis and 1 mL of soil slurry for DNA extraction. After this initial sampling, 2 mL of exudate treatment were added to each microcosm, vortexed, and a second 1 mL aliquot was immediately taken for exometabolomics analysis. At this point, for each microcosm the bottle caps were removed and replaced with a sterile foam stopper for the rest of the experiment to maintain oxic conditions and prevent colonization by contaminating microbes. Microcosms were incubated in an orbital shaker set at 200 rpm at 24°C for 20 days. Each exudate treatment (i.e., HST, HOT and Control) was conducted in triplicate and treatments were added (2 mL) to microcosms on days 0, 1, 2, 3, 4, 5. After day 5, no additional exudate treatments were applied but microcosms were maintained until day 20 which afforded additional samples taken at day 7, 11, and 20 (**Fig. 1C**). Samples were collected at roughly the same time each day and collected with aseptic techniques to ensure no additional microbial influence was introduced. All collected samples were immediately frozen at -80°C until processing.

### Sample Preparation and Extraction for Exometabolomics Analysis

Samples were thawed at 4°C overnight, centrifuged for 20 minutes at 18,000 x g and supernatant was transferred to a pre-weighed 1.5-dram vial and refrozen at -80°C for subsequent lyophilization. Samples were lyophilized for 24 hours until all water was sublimated. Samples were weighed to calculate total exometabolite mass and then were resuspended in 4mL sterile HPLC-grade water. A volume equivalent to 0.50 mg was transferred to a new, pre-weighed 2 mL glass vial and dried under N_2_. Lastly, each sample was resuspended in 500 μL of sterile HPLC-grade water, vortexed for 1 min, and sonicated for 15 min. Two 250 μL aliquots were transferred to new 2 mL vials, respectively, and samples were dried under N_2_. This yielded two 0.25 mg subsamples for analysis by GC-MS and UPLC-MS/MS as described below.

### Targeted UPLC-MS/MS for Phytohormone Analysis of Exometabolites

0.25 mg subsamples were extracted in 75 μL of a spiked methanol solution containing 100% methanol with 65.2 ng/mL ABA-d6, 62.5 ng/mL salicylic acid-d6, and 90.0 ng/mL jasmonic acid-d5 (Sigma). After solvent addition, samples were placed on a shaker plate for 1 hour at the highest speed setting, centrifuged at 3500 x g at 4°C for 5 minutes, and transferred to glass inserts. A final centrifuge step at 3500 x g for 15 minutes at 4°C was completed to ensure any precipitate was in the bulb of the vial insert. Five microliters of exometabolite samples were injected onto a Perkin Elmer UPLC MS/MS system, equipped with a PerkinElmer QSight LX50 Solvent Delivery Module (PerkinElmer). An ACQUITYUPLC T3 column (1 × 100 mm, 1.8 μM; Waters Corporation) was used for chromatographic separation. Mobile phase A consisted of LC-MS grade water with 0.1% formic acid and mobile phase B consisted of 100% acetonitrile. The elution gradient was initially set at 0.1% B for 1 min, which was increased to 55.0% B at 12 min and further increased to 97.0% B at 15 min, then decreased to 0.1% B at 15.5 min. The column was re-equilibrated for 4.5 min for a total run time of 20 min. The flow rate was set to 200 μL/min and the column temperature was maintained at 45°C. Samples were held at 4°C in the autosampler. Detection was performed on a Perkin Elmer QSight™ 220 triple quadrupole MS in selected reaction monitoring (SRM) mode. The transitions monitored for each phytohormone compound can be found in **Table S2**. The MS was operated with ESI voltage 4500 V in positive mode and -3500 V in negative mode. Nebulizer gas flow was set at 350 arbitrary units and drying gas was set to 120 arbitrary units. The source temperature was 315 °C and hot-surface induced desolvation (HSID) temperature was set to 200°C.

### Non-targeted GC-MS Analysis of Exometabolites

Sample preparation was conducted as previously described (29, 57) and as described above for sorghum root exudates. Briefly, dried samples were resuspended in 50 μL of pyridine containing 25 mg/mL methoxyamine hydrochloride (Sigma), centrifuged, incubated at 60°C for 45 minutes, vortexed for 30 seconds, sonicated for 10 minutes, centrifuged briefly, and incubated a second time for 45 mins at 60°C. Samples were cooled to room temperature and centrifuged for 2 minutes. Then, 50 μL of MSTFA + 1% TMCS (Thermo Fisher) was added, samples were vortexed for 30s, centrifuged, and incubated a third time at 60°C for 35 minutes. Samples were cooled to room temperature, centrifuged, and 80μL of supernatant was transferred to glass vial inserts within glass vials. Samples were centrifuged a final time for 10 minutes before analysis. Metabolites were separated with a 30 m TG-5MS column (Thermo Scientific, 0.25mm i.d. 0.25 um film thickness) and detected using a Perkin Elmer Clarus 690 GC coupled to a Clarus SQ 8S mass spectrometer (Perkin Elmer). Samples (1μl) were injected at a 10:1 split ratio onto the column with a 1.0 ml/min helium gas flow rate. The gas chromatography inlet was held at 285°C, and the transfer line was held at 300°C, and the source temp was held at 260°C. The GC oven program started at 80°C for 30s, followed by a ramp of 15°C/min to 330°C, followed by an 8-min hold. Masses between 50–620 *m/z* were scanned at 4 scans/s under electron impact ionization. Injection of QCs were analyzed after every 6th sample to ensure proper instrument function and to detect any analytical variation.

### GC-MS and LC-MS Data Analysis

Non-targeted GC-MS data (both sorghum exudates and exometabolites) were processed within the R statistical software (58) using methods previously described (57). For GC-MS samples, .cdf files were processed through the following workflow: 1) XCMS software was used for preprocessing to identify molecular features (59); 2) feature normalization were further normalized to total ion current (TIC); 3) the package RAMClust (60) was used for clustering features into spectra and prepared for subsequent spectra identification in RAMSearch (61) using external databases including Golm (http://gmd.mpimp-golm.mpg.de/) and NIST (http://www.nist.gov). For exometabolite data, prior to TIC normalization, features were normalized by linearly regressing run order versus QC feature intensities to account for instrument signal intensity drift. For root exudate data, relative quantitation was also normalized by root weight. Targeted quantification of sugars from the GC-MS data was performed using Chromeleon 7.2 (Thermo Fisher). The integrated peak area for each sugar was normalized to its corresponding internal standard with the exceptions that arabinose 13C^5^ was use for xylose, glucose 13C^6^ was used for mannose and sucrose 13C^12^ was used for trehalose and maltose. Quantification was determined using a linear regression of an 8 point standard curve for each sugar. Final concentrations were normalized to root weight. For phytohormone analysis of exometabolites, LC-MS data were processed using Simplicity 3Q (v1.5, Perkin Elmer) bioinformatics software for sample processing. Briefly, the peak area for each phytohormone compound was normalized to internal standard peak area and quantification was assessed using a linear regression against an external calibration curve. Exudate and exometabolite data are provided in **File S1**.

### Soil DNA Extraction and Library Preparation

Total genomic DNA was extracted from the microcosms using the Zymo Quick-DNA Fecal/Soil Microbe Microprep kit. 16S rRNA gene amplicon sequencing was performed on the Illumina MiSeq using 251-bp paired-end reads and the Earth Microbiome Project primers 515F/806R (62), for an average of more than 30,000 reads per sample (see **File S2** for individual sample data). The 16S rRNA partial gene reads were analyzed and reads were demultiplexed using QIIME2 (63) (2019.10). Using DADA2 (64), demultiplexed reads were denoised to produce an amplicon sequence variant (ASV) table and filtered to remove noisy sequences, chimeras and singletons. Feature classification was completed by comparing the ASV table against the trained full-length SILVA classified (silva132.250) database for taxonomic classification. The ASV table was filtered to contain ASVs that were observed in at least 2 samples and the output files were visualized in QIIME2. Samples L_E1_T5_A (HST day 5 rep A), L_E2_T11_A, and L_E2_T20_A (HOT days 11 and 20 rep A) yielded insufficient sequencing results and were excluded from subsequent analyses. The ASV feature table is provided in **File S2**.

### Metagenomics Analysis

Metagenomic DNA from day 0 (control) and day 5 (control and HST) metagenomes (n = 9) was sequenced at the Genomics Shared Resource at the University of Colorado Cancer Center using the NovaSeq6000 platform. Metagenomic DNA from day 20 (control and HST, n = 6) was prepared for metagenomic sequencing using the Nextera XT low input-Illumina library creation kit and samples were sequenced at the Department of Energy Joint Genome Institute on the Illumina NovaSeq 6000. FastQ files were trimmed using Sickle (v1.33) (65). Day 0 and day 5 reads were concatenated within each timepoints/treatments (ie. triplicate controls at day 5) for co-assembly. Day 0 and day 5 (both control and HST) coassemblies, and the day 20 individual assemblies were assembled with IDBA-UD (66). Within each assembly, scaffolds greater than 2.5kb were binned into metagenome-assembled genomes (MAGs) using MetaBAT2 (v2.12.1) (67). MAGs were assessed for completion and contamination using checkM (68). A MAG was retained if it was >48% complete with <10% contamination, and assigned quality following MIMAG guidelines (40). Using dRep (69), MAGs were dereplicated to 99% identity. MAG taxonomy was assigned using GTDB-tk (v1.5.0, R06-RS202) (70). To obtain MAG abundance, trimmed metagenomic reads from individual samples were mapped to the dereplicated MAG set using bbmap (71) (v38.70) at minid=95, and output as sam files which were converted to sorted bam files using samtools (72) (v1.9). CoverM (v0.3.2) was used to determine MAG relative abundance as described in McGivern et al (56). MAGs were annotated using DRAM (73). Biosynthetic gene clusters were detected using the antiSMASH webserver using default parameters (v6.0) (74). See **File S2** for MAG quality, taxonomy, and mapping; **File S3** for DRAM annotations; and **File S4** for raw annotations accessed using DOI: 10.5281/zenodo.5639650.

### Statistical Analyses

Alpha diversity of microbial communities was calculated using the diversity function from the vegan package (75) in R (58) (v4.0.2) using Shannon’s (H), Pielou’s, and richness indices. To estimate beta diversity across treatments, we utilized Bray-Curtis dissimilarity matrix visualized by non-metric multidimensional scaling (NMDS) in R with the ggplot2 package (76) with stress of the non-parametric fit for the goodness of fit for the model. Significance of compositional differences across treatments was quantified using mrpp and the betadisper commands from the vegan package with an ANOVA model. Fold changes for exometabolite dynamics were calculated and converted to log2 abundances via the log function. 2-way ANOVA tests and pairwise comparisons were completed in GraphPad Prism (v8.2.1) using grouped analyses with Sidak’s multiple comparison testing with alpha = 0.05.

## Data Availability

The MAGs, metagenomes, and 16S rRNA amplicon sequencing reads from this dataset have been deposited in National Center for Biotechnology Information under BioProject PRJNA725542 (for biosample accession numbers see **File S2**). MAGs are also provided in archive (https://doi.org/10.5281/zenodo.5142207). Metabolomics (exometabolites and root exudates) data have been deposited to the EMBL-EBI MetaboLights database (pending) (DOI: 10.1093/nar/gkz1019, PMID:31691833) with the identifier MTBLS3164 (77). The complete dataset can be accessed here: https://www.ebi.ac.uk/metabolights/MTBLS3164

## Acknowledgements

A portion of this work was conducted by the U.S. Department of Energy Joint Genome Institute, a DOE Office of Science User Facility, which is supported by the Office of the U.S. Department of Energy under Contract No. DE-AC02-05CH11231 (proposal ID 2049, Wrighton). A portion of this work was performed by the University of Colorado Anschutz’s Genomics Shared Resource supported by the Cancer Center Support Grant (P30CA046934). Sorghum root exudate analyses were conducted under the DOE DER award DE-SC0014395. The authors wish to thank Linxing Yao in the ARC-BIO at Colorado State University for her assistance in analysis of the sorghum root exudates.

